# Corticosteroids elevate intraocular pressure through suppression of TREK-1 signaling

**DOI:** 10.1101/2025.08.14.670390

**Authors:** Sarah N. Redmon, Oleg Yarishkin, Christopher N. Rudzitis, Monika Lakk, Jacques Bertrand, Joseph van Batenburg-Sherwood, Christina M. Nicou, Christopher L. Passaglia, Darryl R. Overby, David Križaj

## Abstract

Clinicians are often forced into the dilemma of whether to battle ocular inflammation or preserve vision imperiled by elevated intraocular pressure (IOP). Anti-inflammatory treatments utilizing glucocorticosteroid regimens may induce glaucoma by chronically elevating IOP via increased trabecular meshwork (TM) resistance to the flow of aqueous humor, but it is not known whether pressure transduction itself is impacted by steroids and how changes in TM mechanosignaling affect conventional outflow resistance and IOP. To address this, we investigated the role of TREK-1 (TWIK-related potassium channel-1), a mechanosensitive K^+^ channel, in regulation of outflow facility, transmembrane signaling and dexamethasone (DEX)-induced ocular hypertension (OHT). The expression of tandem-pore potassium channels in mouse TM cells was dominated by *Trek-1 (Kcnk2*) mRNA, with residual expression of *Traak, Tresk2* and *Twik3* and vanishingly low levels of *Task1 and Trek2*. DEX suppressed *Trek1* transcription by ∼80% but did not affect expression of *Trpv4* and *Piezo1* genes. Chronic DEX administration depolarized the membrane potential of TM cells and elevated IOP in mice whereas the selective TREK-1 agonist ML-402 lowered IOP in rodent OHT models. ML-402 doubled the outflow facility in perfused mouse eyes at all applied pressures and hyperpolarized DEX-treated TM cells. These *in vitro, ex vivo* and *in vivo* results implicate TREK-1 channels in homeostatic regulation of TM mechanosignaling, conventional outflow regulation and IOP homeostasis. Suppression of TREK-1 signaling by corticosteroids underlies OHT and could contribute to steroid glaucoma but this can be obviated by pharmacological stimulation of the channel with cornea-permeant ML-402 eye drops.

## Introduction

Glucocorticoids are widely used in the clinic to combat inflammation associated with autoimmune disorders, eye conditions (Graves’ ophthalmopathy, uveitis), allergies, asthma, heart failure, skin conditions and surgery. Persistent treatment increases the risk for osteoporosis, weight gain, high blood pressure and cognitive defects (*1*) and may cause permanent vision loss due to chronic elevation of intraocular pressure (IOP) (*2, 3*). Clinicians are thus often forced into the dilemma of whether to battle ocular inflammation or preserve vision imperiled by elevated intraocular pressure (IOP); an estimated one-third of the population is moderately steroid responsive (rise of IOP 6–15□mmHg from baseline with steroid treatment) and 4–6% of the population highly steroid responsive (IOP increase of >15□mmHg from baseline or IOP over 31□mmHg after steroid exposure) (1 - 3). Steroid sensitivity is augmented in patients with primary open angle glaucoma (POAG), 90% of whom display elevated blood cortisol levels and glucocorticoid-induced ocular hypertension (OHT) (2, 3).

Steroids elevate IOP by augmenting the resistance of the conventional outflow pathway to the flow of aqueous humor (AH) (*4*). The pathway is comprised of episcleral vessels, the inner wall endothelium of the canal of Schlemm and trabecular meshwork (TM) comprised of cells that populate three distinct ECM layers, are highly mechanosensitive (*5-7*), contractile (*8, 9*) and may exhibit properties of a pressure-activated biomechanical pump (*10*). Rabbit, dog, rodent, non-human primate and human TM respond to glucocorticoids with changes in cell-cell adhesion, cell cycle, metabolism, inflammatory pathways, TGFβ2 signaling and expression of cytoskeletal, extracellular matrix (ECM) proteins (*11-18*). We don’t, however, know whether steroid-associated signaling causes OHT or represents its epiphenomenon and it would be useful to determine whether and how steroids affect IOP sensing and homeostasis and conversely, whether manipulation of TM mechanosignaling impacts steroid-induced OHT.

In this study, we investigated the biological function and steroid-susceptibility of TREK-1, a member of the tandem-pore potassium (K_2P_) channel family encoded by the KCNK2 gene. TREK-1 is a *bona fide* mechanochannel that is sensitive to stretch, temperature, pH and C22:6 ω-3/C18:2 ω-6 polyunsaturated fatty acids (*19-21*). TREK-1 channels subserve the K^+^ leak conductance in human TM cells, regulate the permeability of TM monolayers, can be activated by modest (<15 mm Hg) pressure steps (*5, 20, 22*) and are affected by primary open angle glaucoma (POAG) (*23, 24*). Studies in heart, vasculature, gut, and bladder preparations associated TREK-1 -mediated K^+^ efflux with protection from mechanical stress and inflammation (*25-27*). ECM interactions were shown to suppress TREK-1 activity in TM cells isolated from POAG patients (*23*) but it remains unclear whether TREK-1 activity contributes to modulation of conventional outflow in a manner that could lead to OHT.

## Materials and Methods

### Animals and ethical approval

C57BL/6J mice aged 1.5-4 months were purchased from JAX (Bar Harbor, ME) and housed at the Moran Eye Center vivarium in individually ventilated enclosures at 21 °C with a 12 h light-dark cycle and *ad libitum* food and water. Rats were purchased from Envigo (Indianapolis, IN) and housed in the University of South Florida vivarium under similar conditions. Data were gathered from 1 male and female animals with no noted gender differences. The experiments were performed in compliance with institutional IACUCs (Universities of Utah and Miami) and the ARVO Statement for the Use of Animals in Ophthalmic and Vision Research.

### Reagents

The TREK-1 agonist was synthesized as described by Lolicato et al. (*28*) and stored at -80° C. 20-40 mM solutions were obtained by diluting aliquots of the stock solution before each experiment. Salts were purchased from Sigma (St. Louis, MO) or VWR (Radnor, PA).

### mTM/pTM purification and culture

Primary human TM cells (pTM) were isolated according to consensus recommendations (*29*) from human donor eyes (Lions Eye Bank at the Moran Eye Center) (*9, 30, 31*) and used in concordance with the tenets of the WMA Declaration of Helsinki and the Department of Health and Human Services Belmont Report. The trabecular meshwork was dissected out of the anterior eye, plated on 10 mg/mL collagen I coated tissue flasks with cells migrating from the tissue over 2 - 4 weeks. Trypsinized cells were replated on collagen-I coated flasks in the TMCM medium (ScienCell, Carlsbad, CA) supplemented with growth serum (TCGS, ScienCell), 2% FBS, 100 U/ml penicillin and 100 mg/mL streptomycin. The experiments were conducted with P1-P4 cells, which were validated as described (*7, 31, 32*).

The isolation of mTM cells followed the protocol in (*33*). Briefly, retina, choroid, vitreous, lens and cornea were removed, limbal rings dissected, rinsed in PBS containing 1% AA (ThermoFisher) and digested (1% AA, 4 mg/mL collagenase A (MilliporeSigma), 4 mg/mL BSA in PBS) for 2 hours at 37°C. Dissociated cells were transferred to flasks at 37°C and incubated for 24 hours. 0.001% magnetic polystyrene smooth surfaced beads (2.0-2.9 µm, Spherotech Inc, Green Oaks, IL) were added, and cells detached with TrypLE Express™ (GIBCO; ThermoFisher Scientific, Waltham, MA). Phagocytosing cells were magnetoprecipitated, re-suspended in 6 mL of media and seeded into DMEM media (1 g/L glucose, GIBCO) supplemented with 10% FBS (Genesee. El Cajon, CA) and 1% AA (ThermoFisher) at 37°C and 5% CO_2_. The cells were used between passages 2-4. TM identity was validated by the expression of *Myoc, Chi3L, Acta2* and *Timp3* marker genes and DEX-induced upregulation of *Myoc* (*33*).

### Gene expression analysis

mTM cells were used for analyses of mRNA expression. Total RNA was isolated with the Arcturus RNeasy Plus Micro Kit (Qiagen) and 500 ng of total RNA used for reverse transcription. First-strand cDNA synthesis and PCR amplification of cDNA was performed using qScript™ Ultra SuperMix cDNA synthesis kit. SYBR Green based real-time PCR was performed using Apex qPCR Master Mix (Genesee Scientific). The results were performed in triplicate of at least three experiments. The comparative C_T_ method (ΔΔC_T_) was used to measure relative gene expression where the fold enrichment was calculated as: 2^-[ΔCT (sample) - ΔCT (calibrator)]^ after normalization. *Gapdh* levels were utilized to normalize fluorescence signals. Primer sequences are shown in Table 1.

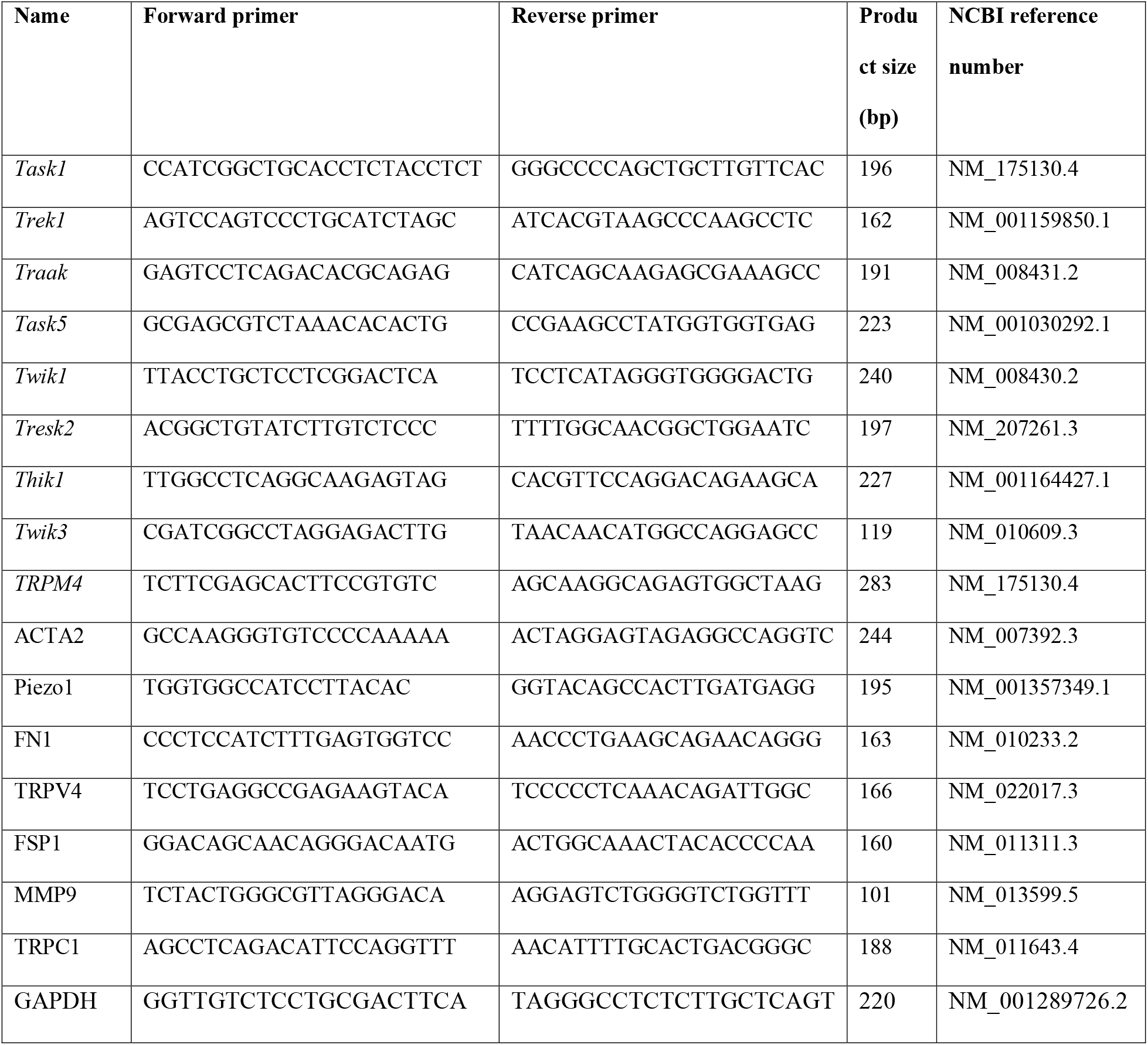

### Immunohistochemistry

Mice were euthanized, and eyes fixed in 4% PFA for one hour at room temperature (RT) (*34, 35*). The anterior eye tissue was cryoprotected in 15% sucrose and 30% sucrose for 45 min at room temperature and overnight at 4°C, respectively, and the eyes embedded in OCT (Tissue-Tek Sakura, Torrance, CA). 12 µm slices were cut with a cryostat, mounted on Superfrost plus slides (ThermoFisher), dried and kept at -80°C. For staining, cryosections were rinsed in PBS (3 × 5 min) and blocked in 5% FBS and 0.3% Triton-X in 1X PBS for 30 min (sections) at RT. Primary antibodies (mouse αSMA 1:500; Abcam; rabbit TREK-1, 1:200, Santa Cruz Biotechnology, Santa Cruz, CA) were prepared in 2% BSA and 0.2% Triton-X in 1X PBS. Cryosections were incubated overnight at 4°C, rinsed 3 times for ∼5 minutes in 1X PBS, incubated with secondary antibodies for 1hr at RT and washed 3x for 5 min in 1X PBS. Secondary antibodies were goat anti-mouse Alexa Fluor 594 1:1000 Invitrogen; goat anti-rabbit Alexa Fluor 488 or 594 1:1000 (ThermoFisher Scientific; goat anti-guinea pig Cy3 1:1000 Jackson ImmunoResearch). Cryosections were counterstained (DAPI-Fluoromount-G, Southern Biotech, Birmingham AL) and coverslipped. Immunofluorescence was acquired with confocal microscope (Olympus FV1200, 1024 × 1024 pixels with a 20x water superobjective).

### Electrophysiology

Whole-cell patch current was recorded (*20, 31, 36*) at room temperature (22 – 24°C) with borosilicate pipettes (5 - 8 MΩ; WPI, Sarasota, FL) pulled on the P-2000 puller (Sutter Instruments, Novato CA). Pipettes were fabricated from borosilicate glass capillaries (WPI; outer diameter 1.5 mm, inner diameter 0.84 mm). The external solution contained (in mM): 126 NaCl, 2.5 KCl, 1.5 MgCl_2_, 1.5 CaCl_2_, 1.25 NaH_2_PO_4_, 26 NaHCO_3_, 10 D-glucose (pH 7.4, adjusted with NaOH). The pipette solution contained (in mM): 135 potassium D-gluconate, 10 KCl, 1 MgCl_2_, 10 HEPES, 0.5 EGTA, 0.01 CaCl_2_, 0.3 Na-GTP (pH 7.3, adjusted with KOH) and 1 Mg-ATP. Acquisition was controlled via a Multiclamp 700B amplifier, Digidata 1550 board and Clampex 10.7 acquisition software (all Molecular Devices; Danaher Medical Technologies, San Jose, CA). Cells were held at the resting potential (typically between -30 mV and -40 mV), with currents were sampled at 10 kHz, filtered at 5 kHz with an 8-pole Bessel filter, and analyzed with Clampfit 10.7 (Molecular Devices) and OriginPro 8 (Origin Lab).

### iPerfusion flow measurement

Outflow facility was measured in enucleated eyes from C57BL/6J mice (N = 6 mice; 11-week-old males; Charles River UK Ltd., Margate, UK) using *iPerfusion (31, 37, 38*). Briefly, enucleated eyes were affixed to a support platform and submerged in a PBS bath at 35°C. The anterior chamber was cannulated using a glass micropipette pulled to a 100 µm diameter beveled tip. The perfusion fluid was Dulbecco’s PBS containing divalent cations and sterile (0.25 µm) 5.5 mM glucose. Ipsilateral eye was perfused with ML-402 (30 µM) and the contralateral eye perfused with vehicle. IOP was set to 9 mmHg for 1 hour to pressurize and acclimatize the eye to the perfusion environment and to allow sufficient time for the TREK-1 agonist to reach the anterior chamber. Flow was measured over 8 increasing pressure steps from 6.5 to 17 mmHg. Steady state for each step was evaluated when the ratio of the flow rate to pressure changed by less than 0.1 nl/min/mmHg per minute over a 5-minute window (Sherwood et al., 2016). The stable pressure, *P*, and flow rate, *Q*, were calculated over the last 4 minutes of each step, and a power-law relationship of the form

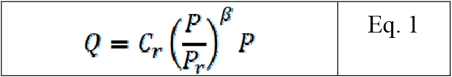

was fit to the *P* − *Q* data. The reference outflow facility, *C*_*r*_, represents the value of outflow facility at a reference pressure *P*_*r*_ of 8 mmHg, and *β* characterizes the non-linearity of the *Q* − *P* relationship (Sherwood et al. 2016). The relative difference in *C*_*r*_ between treated and untreated contralateral eyes was calculated as the ratio of *C*_*r*_ in the treated eye relative to that in the contralateral control eye minus unity. Facility values and relative changes in facility are reported in terms of the geometric mean and the 95% confidence interval on the mean.

### Tonometry and IOP elevation

#### DEX-induced OHT

Mice were habituated to handling, with topical 0.1% DEX sodium phosphate ophthalmic eyedrops (Bausch & Lomb Inc.) administered to awake animals three times a day (10-11 AM, 2-3 PM, 6-7 PM) for ∼12 weeks and age-matched cohorts treated with sterile PBS drops. The animals were gently restrained for ∼30 seconds after drop application to allow for glucocorticoid permeation. IOP in Avertin-treated mice was measured with a rebound tonometer (Tonolab; Colonial Supplies, Charleston SC) between noon and 2 PM, with each IOP data point representing an average of 10 to 20 tonometer readings. No sex differences were noted, and data from male and female animals was pooled.

#### IOP telemetry

Rats were anesthetized, a fluid-filled silicone microcannula was implanted in the anterior chamber of one eye, and the cannula was connected to a wireless pressure sensor via a plastic coupler secured to the skull (*39, 40*). When the animal woke, IOP was transmitted at 0.25Hz round-the-clock to a laptop for display and storage. At sporadic times, a drop of drug (ML-402; 40 µM) or vehicle (PBS saline) was instilled on the implanted eye.

### Statistical Analysis

Group sizes were determined based on preliminary experiments, with data analyzed by GraphPad Prism 10.0. Two-way repeated measures ANOVA was for analyses of longitudinal IOP data in ipsi/contralateral eyes treated with vehicle PBS and ML-402. Paired t-test was utilized to compare two groups with pre-injection and post-injection. The details are included in the figure legends for each panel.

## Results

### Histology: TREK-1 channel expression in the anterior eye of the mouse

TREK-1 expression in anterior segment of eyes from wild-type C57BL/6 mice was studied by antibody labeling. Immunoreactivity (TREK1-ir) was observed in nonpigmented cells of the ciliary body (CB), corneal epithelium and the trabecular meshwork (TM) (Fig. 1A), where it was confirmed by colocalization with α-smooth muscle actin (αSMA). Relative expression of tandem-pore potassium (K2P) family gene isoforms was ascertained in mouse TM (mTM) cells that manifest the canonical transcriptional signature indicated by robust expression of *Myoc* (myocilin), *Mgp* (matrix Gla protein), *Chil3l1* (Chitinase-3-like protein1) and *Mmp2* (matrix metalloproteinase 2) (*33*). The mTM K2P transcriptome was notable for the prevalence of *Trek-1* (TWIK-related K□ channel, encoded by the *Kcnk2* gene) transcripts, modest expression of *Traak* (TWIK-related arachidonic acid-stimulated K□ channel, *Kcnk4*) and *Thik-2* (tandem-pore-domain halothane-inhibited K□ channel, *Kcnk12*) and low to undetectable expression of *Trek-2* (*Kcnk10*; a mechanosensitive paralog of Trek-1), *Twik-3* (TWIK-related acid-sensitive K□ channel, *Kcnk9*), *Task1* (TWIK-related acid-sensitive K□ channel, *Kcnk3*) and *Task-5* (TWIK-related spinal cord K□ channel; *Kcnk15*) genes. The overall K2P channel expression pattern in mTMs is thus similar yet distinct from primary human TM (pTM) cells (*20*).

**Figure 1.**
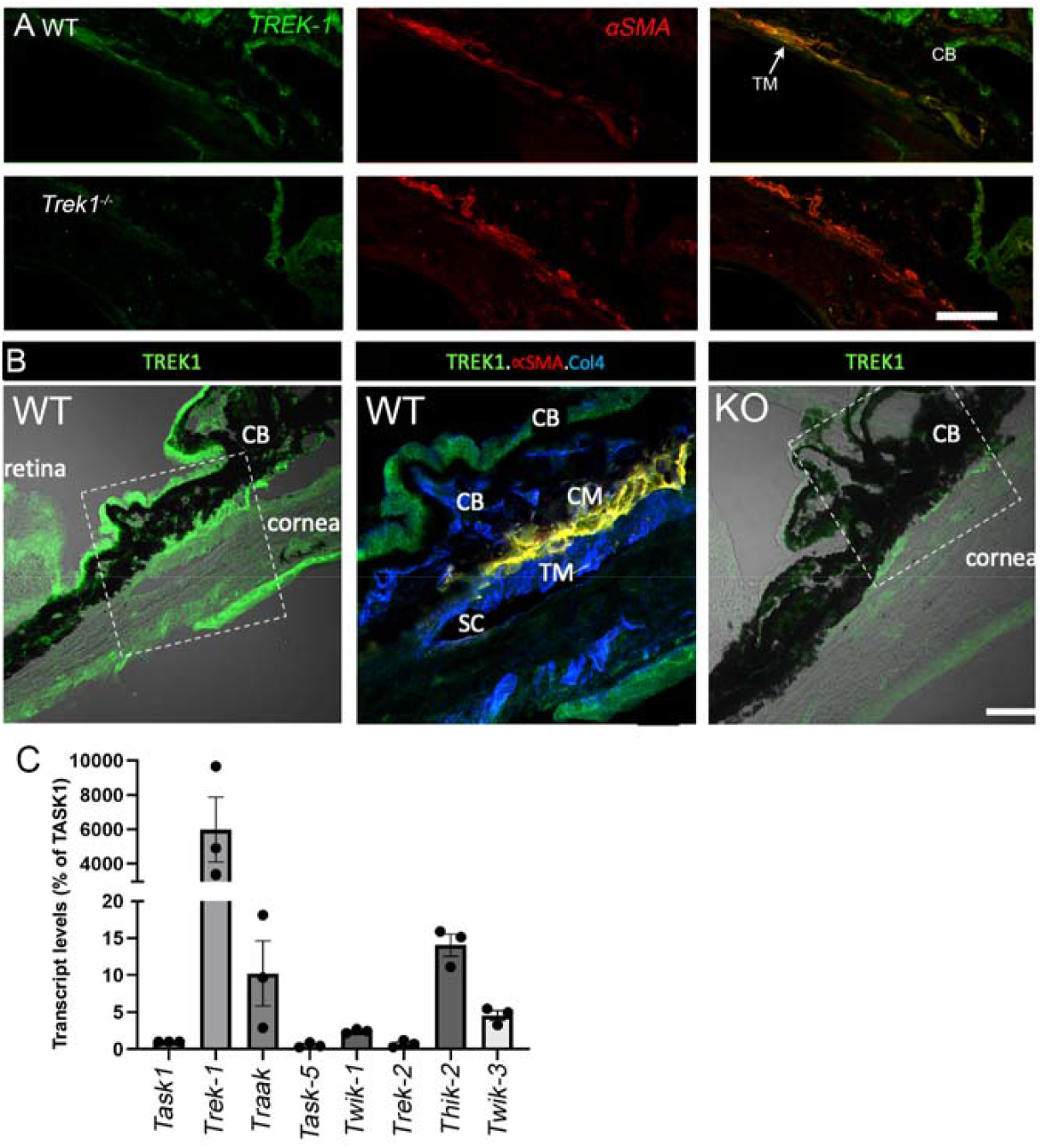
TREK-1 expression in the anterior eye, C57BL/6 mice. (A) TREK-1-ir colocalizes with αSMA in the TM and labels epithelial layers of the cornea and the ciliary body (CB). (B) Lower-magnitude view, the dotted quadrangle denotes the TM. TREK-1-ir colocalizes with αSMA and Col4 (Collagen IV)-ir, with additional expression in corneal epithelium and the CB. Scale bar = 50 µm. (C) qPCR, mTM cells. Relative K2P gene expression is dominated by the *Trek-1* gene. Error bars, ± S.E.M.

### Pharmacological TREK-1 activation doubles the *ex vivo* outflow facility

TREK-1 channels subserve the resting K^+^ leak conductance in pTM cells and modulate the permeability of pTM monolayers (*20*) but their potential roles in outflow modulation and IOP homeostasis remain unclear. We utilized *iPerfusion*, a cannulation-based fluidic system that quantifies the facility via differential pressure measurements (*37, 38*) to test the effect of ML-402, a selective TREK channel activator (*28*), on conventional outflow in mouse eyes. The eyes were pressurized at 8 mm Hg and subjected to incremental pressure steps with and without the agonist (30 µM). ML-402 approximately doubled the flow rate through the outflow pathway at each level of applied pressure. *C*_*r*_ for ML-402 - treated eyes 7.9 [6.3, 9.9] nl/min/mmHg vs. 3.4 [2.4, 4.9] nl/min/mmHg for vehicle-treated eyes (geometric mean [95% CI]; Fig. 2), corresponding to an average increase of 125 [86, 172]% relative to contralateral vehicle-perfused eyes (*P* = 0.0006, n = 6) (Fig. 2C). These findings establish TREK-1 activity as a positive regulator of conventional outflow.

**Figure 2.**
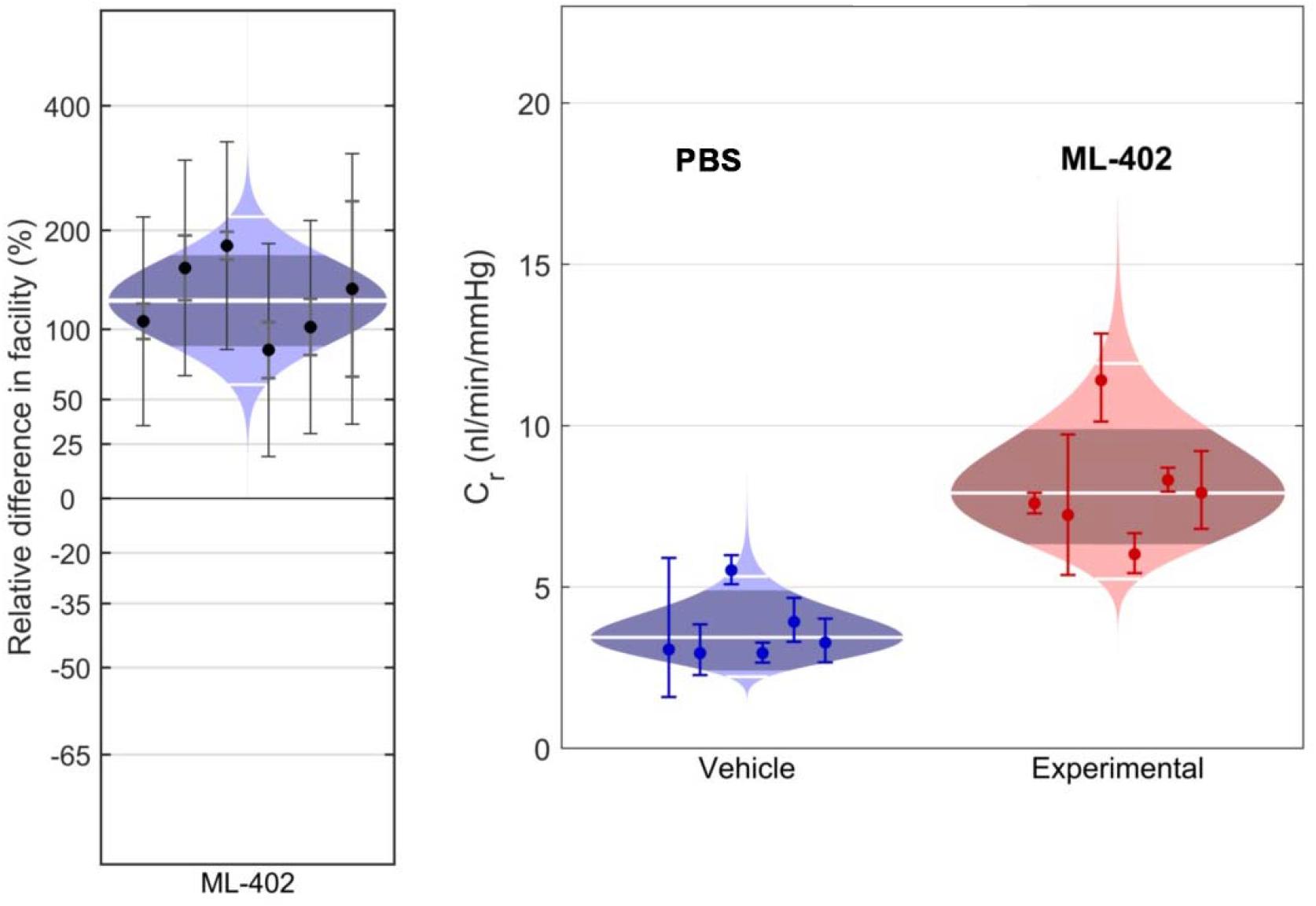
TREK-1 agonism increases outflow facility in enucleated mouse eyes perfused with ML-402 (30 µM) versus vehicle (D-PBS + 5 mM glucose). (A) Cello plot showing the relative difference in reference outflow facility (*C*_*r*_) between contralateral eyes perfused with 30 µM ML-402 vs. vehicle, showing an average increase in *C*_*r*_ of 125 [86, 172]% (N = 6). (B) Facility values for vehicle and ML-402-perfused eyes. Error bars on each data point correspond to 95% CI for each measurement (C) Control facility plotted against experimental cohort. The central white lines represents the geometric means, outer white lines represent 95% data boundaries, and central dark regions the 95% CI.

### TREK-1 activation lowers IOP in an acutely elevated mechanical model

The speed of TREK-1 activation by mechanical stimuli (msec) (*19, 21*) suggests that the agonist could influence IOP at temporal dynamics beyond the limits of perfusion-based systems. Given that mouse eyes are too small for implantation with wireless IOP telemetry systems, we implemented a telemetry protocol in the Brown-Norway rat model (*39, 40*). Proof-of-principle experiments (N = 3/3) showed that corneal administration of the TREK-1 agonist ML-402 (40 µM) lowered IOP with elevated IOP (P < 0.05) (Fig. 3).

**Figure 3.**
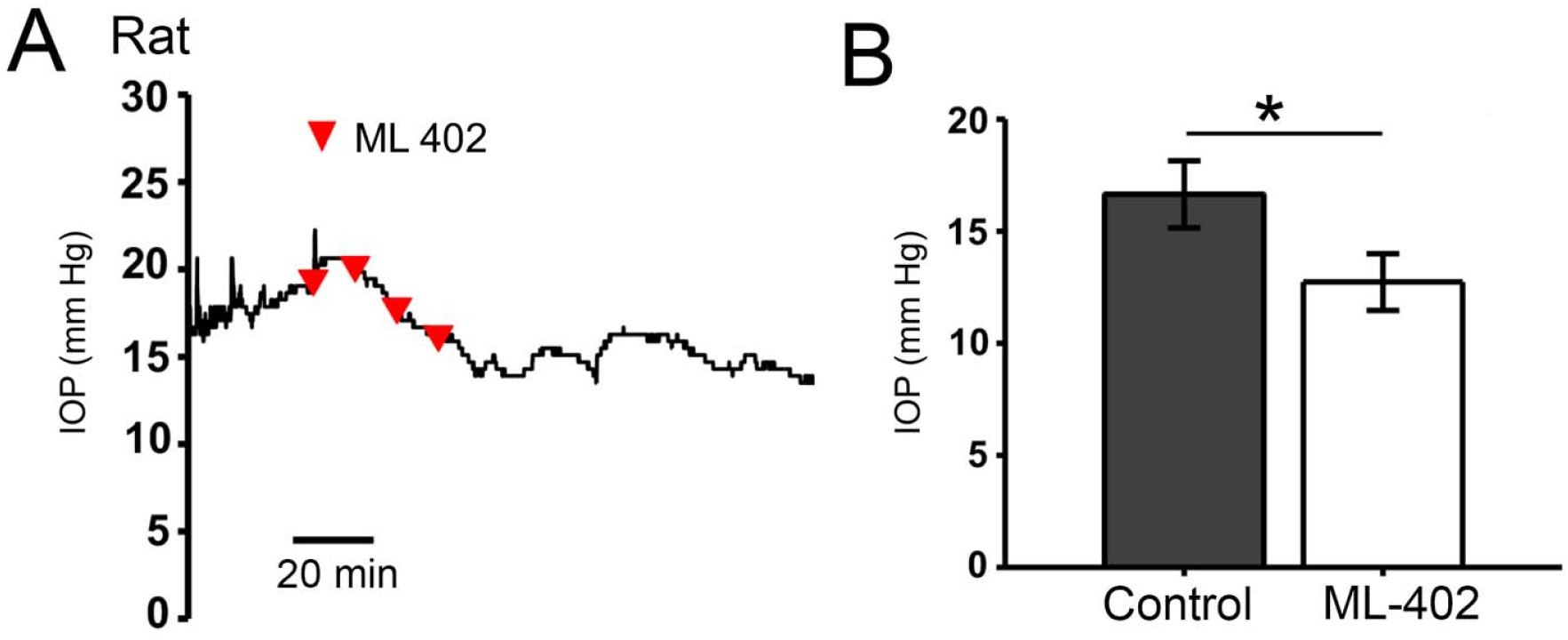
Acute IOP elevations in mice. (A & B) IOP was elevated by adjusting the height of the saline-containing reservoir connected to cannulated eyes. (A) Steady-state IOP is unaffected by PBS drops whereas (B) ML-402 (40 µM) eye drops induce IOP lowering (n = 3, N = 3). Mean ± S.E.M. (C) Rat telemetry. Representative recording of IOP lowering induced by ML-402 in rat eyes. Paired sample t-test; shown as means ± SEM; *, P < 0.05.

### TREK-1 activation lowers IOP in glucocorticoid model of OHT

We employed tonometry to test whether TREK-1 activation modulates IOP in hypertensive eyes that experienced chronic exposure to glucocorticosteroids. As shown previously (*14*), daily dexamethasone (DEX) (0.1%) administration induced gradual increase in IOP that stabilized at 19.4 ± 0.3 mm Hg at ∼5 Weeks. A single eye drop of ML-402 (40 µM) was sufficient to reduce IOP for 4 -6 hours (N = 11; P < 0.0001) (Fig. 4), suggesting that induced TREK-1 activity could protect eyes challenged by steroid overexposure.

**Figure 4.**
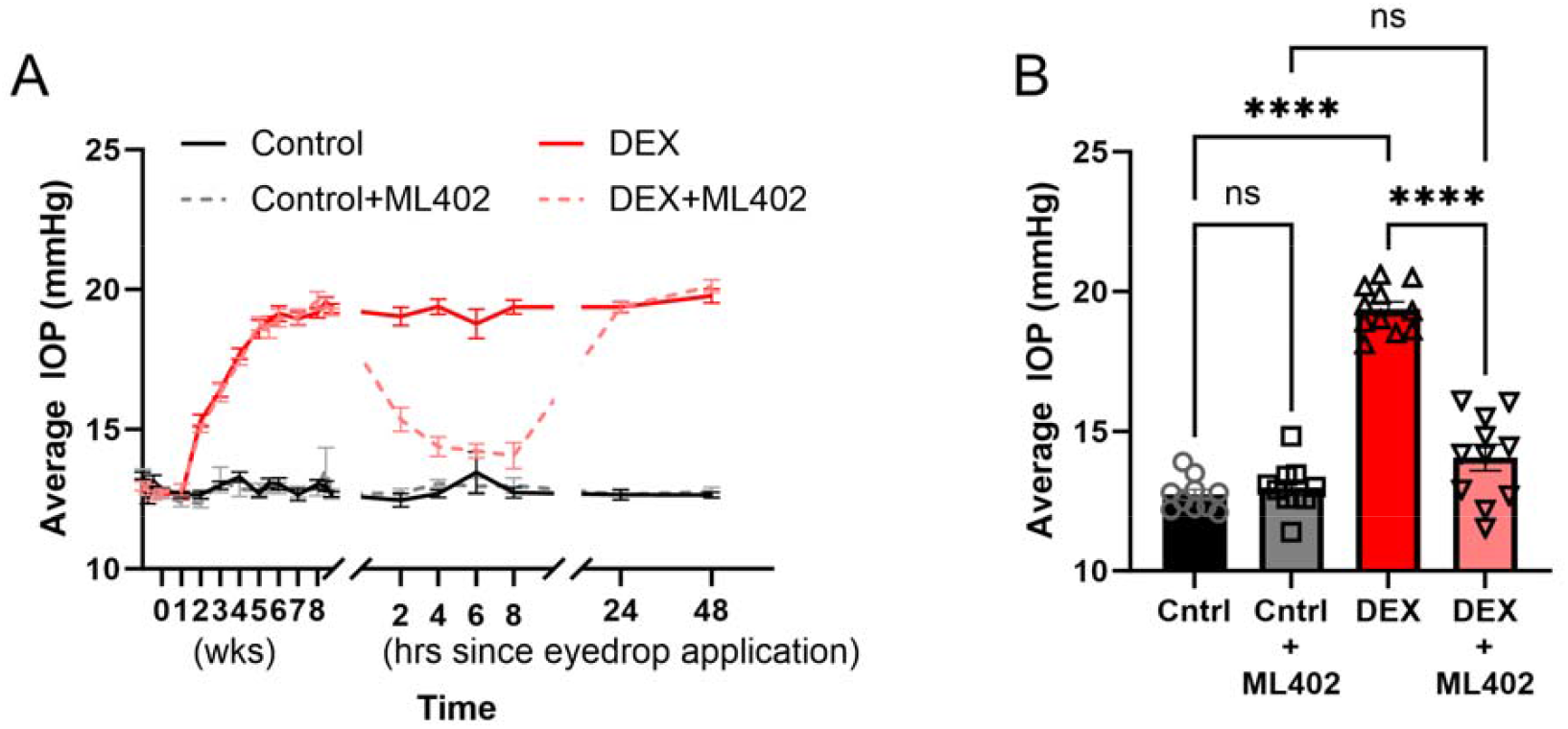
TREK-1 activation lowers IOP in mouse eyes exposed to chronic DEX treatment. (A) Time course of IOP elevations in eyes topically treated with DEX (3x daily) and vehicle (PBS) in the presence and absence of ML-402. ML-402 (N = 11; pink trace) but not PBS (N = 11; red trace) lower IOP in DEX-treated eyes. Neither PBS (N = 10) nor ML-402 (N = 10) affected normotension (gray and black traces). (B) Data summary for the experiments shown in A, 8 hours after ML-402 treatment. Repeated-measures one-way ANOVA with Dunnett’s post-hoc comparisons, presented as average mean ± SEM. ****, P < 0.001.

### DEX suppresses constitutive TREK-1 activity, an effect partially mitigated by pharmacological TREK-1 activation

To ascertain whether the DEX affects TREK-1 activity, we conducted whole cell recordings of resting and evoked transmembrane currents in pTM cells from donors without history of glaucoma. As reported previously (*20, 22, 36, 41*), membrane potentials in current-clamped cells ranged between -30 and -40 mV (Fig. 5A). Consistent with TREK-1 activation, the cells were hyperpolarized by ML-402 (40 µM) (n = 8). DEX treatment (500 nM, 7 days) depolarized the cells’ resting potential to ∼ -16 mV and attenuated the amplitude of ML-402-evoked hyperpolarizations (n = 6; P < 0.05). Hence, glucocorticoids reduce the steady-state leak K^+^ conductance together with the size of the ‘reserve’ pool of activatable TREK-1 channels.

**Figure 5.**
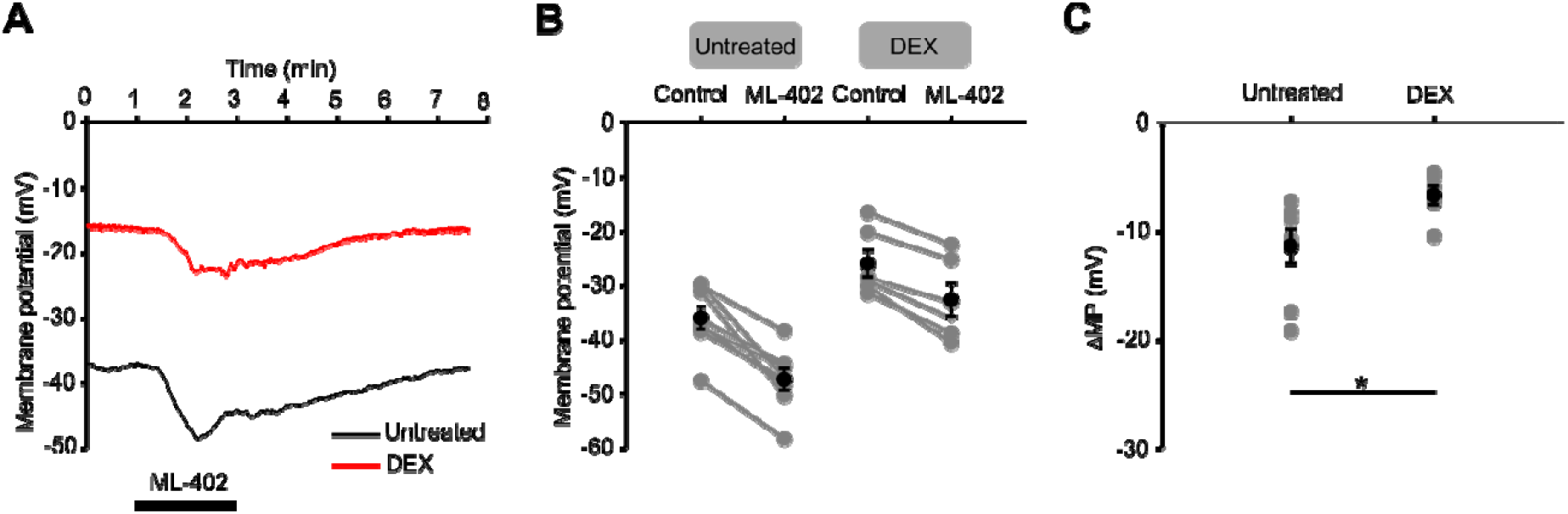
DEX depolarizes pTM cells and reduces the amplitude of the ML-402-evoked current. Whole cell recording, current clamp. (A) Representative recordings of Vrest and ML 402-evoked current from control and DEX-treated cells. (B) Summary of Vrest and ML-402-evoked membrane potentials in control (n = 8) and DEX-treated (n = 6) preparations and (C) The amplitude of the TREK-1 current. Steroid treatment attenuates the ML-402-evoked current. Black markers indicate means ± SEM, paired-sample t-test, * P < 0.05.

### Glucocorticoid treatment downregulates KCNK2 expression and modulates expression of genes that encoding K2P and myofibroblast marker genes

The simplest explanation for reduced functional expression of TREK-1 in DEX-treated cells (Fig. 5) and its role in OHT (Fig. 3) is that glucocorticoids modulate the expression of Kcnk2 and/or other genes involved in TM mechanosensing. As reported previously for pTM cells (*42*), DEX upregulated mTM expression of *Fsp1* (Fibroblast Specific Protein-1), a TGFβ2-sensitive myofibroblast marker gene activated downstream from RhoA. We also observed suppressed expression of *Trpm4, Acta2* and *Mmp9* (Fig. 6) and ∼80% downregulation of *Kcnk2* expression (P < 0.5, N = 4) whereas expression of *Piezo1, Trpv4* or *Trpc1* genes appeared unaffected by steroid exposure.

**Figure 6.**
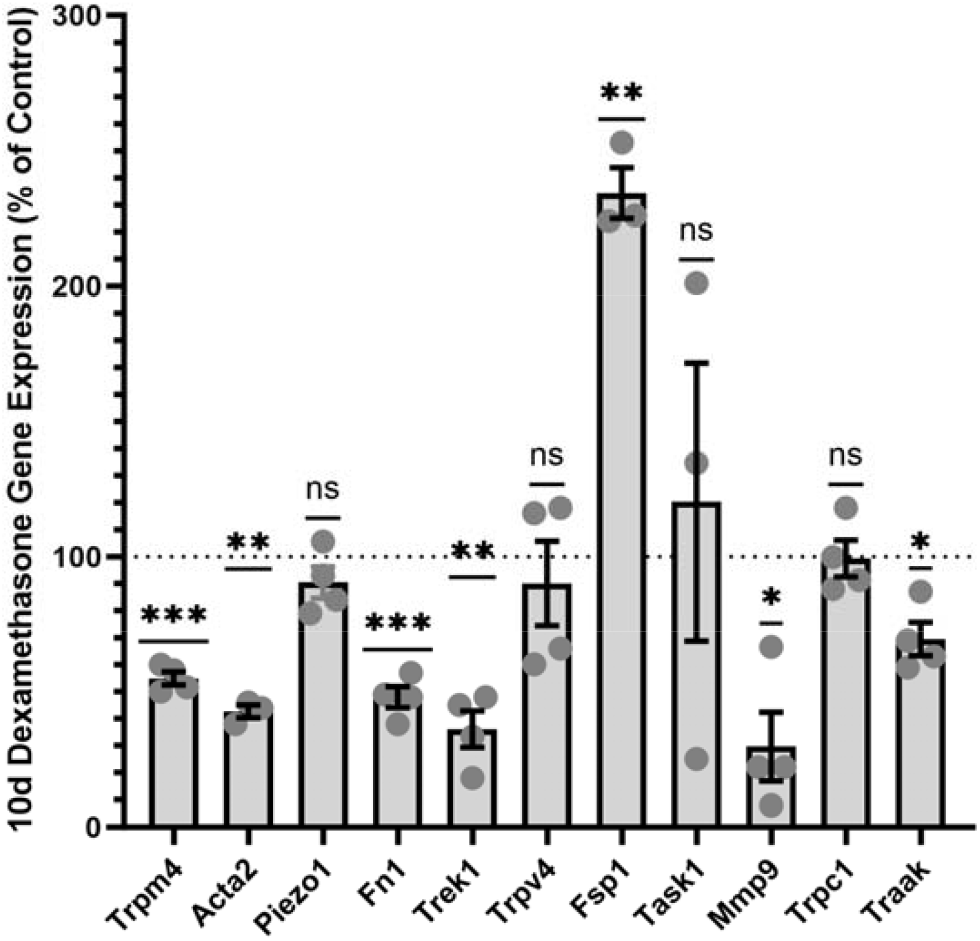
DEX modulates expression of K2P and myofibroblast marker genes. mTM, 10-day exposure to DEX (100 nM), semiquantitative qPCR. ∼fold change in relative abundance of transcripts that encode fibrosis- and mechanosensation-related proteins (N = 4) (*Trpm4*, transient receptor potential melastatin 4; *Fn1*, fibronectin 1; Fsp1, fibrosis-related gene isoform 1; *Mmp9*, matrix metalloproteinase isoform 9; *Trpc1*, transient receptor potential canonical isoform 1). DEX downregulates *Trek-1* expression without impacting the expression of *Trpc1, Task1, Piezo1* and *Trpv4* genes. One-way ANOVA, mean ± SEM. *, P < 0.05; ** P < 0.01; ***P < 0.001

## Discussion

This study brings insights into the molecular mechanisms that subserve pressure-dependent IOP homeostasis in normo- and hypertensive eyes and explores the interaction between steroid overexposure and dysregulated pressure sensing within the TM. While glucocorticoid therapy is commonly used to treat many ocular and systemic conditions, its many serious health risks include disruption of the HPA axis, impaired cardiovascular function and blindness: ∼40% of people without glaucoma develop steroid-induced OHT, with glaucoma risk rising to ∼90% in POAG patients and ∼80% in patients with Cushing’s Syndrome (*1-3*). Increase glucocorticoid levels during aging are likely to additionally predispose individuals towards IOP dysregulation (*43*). Our study has four major sets of findings that bring insight into the molecular mechanism of steroid-induced OHT. First, we found that TREK-1 channels facilitate conventional outflow, presumably contributing to IOP stability in healthy eyes (*44*). Second, TREK-1 activation reduces acute and chronic OHT. Third, exposure to DEX depolarized TM cells and elevated IOP, effects that correlated with downregulation of *Trek-1* gene and TREK-1 current. Interestingly, expression of *Piezo1* or *Trpv4* genes appeared unaffected by the steroid. Finally, glucocorticoid-induced OHT was reversed by a selective TREK-1 agonist (*28*).

The similarities in the anatomy and physiology of the outflow pathway in mice and humans (*29, 45*), conserved expression patterns for phenotype-defining genes (e.g., MYOC, PITX2, ACTA2, TRPV4, CHI31L, TIMP3, MMP9), similar glucocorticoid sensitivities of mouse and human TM (*33, 46-48*) and comparable stiffnesses between DEX-treated mouse and glaucomatous human outflow tissues (∼69 kPa) (*47, 49*) have long recommended utilization of mouse models of DEX-induced OHT for gaining insight into the disease. IOP measurements in mice have been historically complemented by *in vitro* experiments in human TM cells, with few if any, studies having attempted to correlate *in vivo* tonometry in mice with *ex vivo* analyses of outflow facility and *in vitro* electrophysiological and molecular experiments in the same species. The mTM response to DEX: transcriptional suppression of *Mmp9*, which encodes a Type IV collagenase necessary for maintenance of normotension (*50*) and upregulation of *Fsp1*, a myofibroblast marker gene linked to RhoA-dependent cytoskeletal remodeling (*42*) mirror fibrosis, suppressed ECM turnover and increased stiffness reported for glucocorticoid-treated human TM cells (*15, 47, 49, 51-53*). Expression of genes that encode mechanosensitive channels is similarly translatable, with TM of both species exhibiting prominent expression of *KCNK2/Kcnk2, TRPV4/Trpv4, PIEZO1/Piezo1* and *TRPC1/Trpc1* but not PIEZO2/*Piezo2* genes (*9, 33*) (Fig. 6). Expression of cognate *Trek-2* mRNA was low in human and mouse preparations (*20*) whereas mTM but not pTM showed modest expression of *Traak* (which encodes another mechanosensitive channel) (*25*) and *Thik-2* (which encodes a ‘nonfunctional’ K2P subunit) (*54*). Unlike pTM cells (*20*), mTMs showed vanishingly low expression of *Task-1*. Overall, these data suggest that mechanosensitive K^+^ efflux in mTMs is carried by TREK-1, with residual involvement of homomeric TRAAK channels and/or TREK-1-TRAAK heteromers.

We utilized iPerfusion, which obviates pressure-independent contributions from inflow (ciliary body), choroid (uveovortical outflow) and suprachoroidal (uveoscleral outflow) pathways (*37*), to evaluate the TREK-1 dependence of the conventional outflow pathway. Consistent with prominent *Kcnk2* expression and pressure-dependence of TREK-1 (*20*), exposure to ML-402 approximately doubled the facility at every applied pressure (Fig. 2). This result supports the hypothesis (*20*) that TREK-1 channels partition between constitutively active channels that subserve V_rest_, and ‘reserve’ channels that remain available for activation by agonists and/or pressure. Precedents from GI tract, bladder, uterus, lung and cardiovascular smooth muscle cells, epithelial and endothelial studies (*25, 26*) are consistent with the conclusion that TREK-1 activation promotes tissue permeability via hyperpolarization, [Ca^2+^]_i_ lowering and cytoskeletal relaxation. The relative contribution of trabecular vs. Schlemm’s canal vs. episcleral TREK-1 pools to ML-402-induced facility increases and IOP lowering needs to be investigated in future studies relying on conditional *Kcnk2* ablation. The link between K^+^ efflux and attendant hyperpolarization/relaxation of the TM is supported by facility increases and IOP lowering in mouse, rat and pig preparations treated with pharmacological activators of K_ATP_ (Kir 6.1) and BK_Ca_ (KCa1/.1) channels (*55-59*). We cannot exclude involvement of a parallel, non-mutually exclusive osmotic redistribution mechanism in which paracellular aqueous fluid flow is facilitated by the reduction in tortuosity and expansion of extracellular volume fraction resulting from the loss of intracellular K^+^ and aquaporin-mediated water efflux (*60, 61*). Taking into account that these effects are antagonized by TRPV4, a swelling-activated nonselective cation channel, which elevates [Ca^2+^]_i_, promotes cell swelling and increases actomyosin contractility to reduce the (*in vitro*) facility and elevate IOP (*33, 34, 41, 61*), it is plausible that TREK-1 helps maintain the homeostatic IOP setpoint by serving as a brake for pressure- and TRPV4-mediated depolarization, Ca^2+^-dependent Rho signaling and cytoskeletal dynamics (*7, 9, 22, 33, 34, 62*). Aging, mechanical stress, chemical factors (e.g., glucocorticoids and TGFβ2) and circadian factors could modulate the setpoint by controlling the relative expression, trafficking and activation/inactivation states of mechanosensitive TRP, Piezo and K2P channels. For example, TGFβ2 may elevate IOP by promoting the expression of TRPV4 (*9*) whereas glucocorticoids elevate IOP through suppression of TREK-1 transcription (Fig. 5). Interestingly, TRPV4 and TREK-1 are thermochannels with comparable temperature activation profiles, sensitivities to arachidonic acid, phosphatidyl inositol biphosphate, cholesterol, protons and participation in cell-ECM interactions (*22, 24, 26, 30, 62-64*), with *Trek1*^*-/-*^ mice showing increased sensitivity to mechanical stressors (mechanical allodynia and hyperalgesia) (*65, 66*) while *Trpv4*^*-/-*^ mice exhibit decreased mechanical hyperalgesia (*67*). We propose that the dysregulated homeostatic setpoint indicated by OHT can be restored through TRPV4 inhibition (*9, 33*) or TREK-1 activation (Figures 3 and 4).

In summary, given that clinicians can be forced into the dilemma of whether to battle ocular inflammation or preserve vision imperiled by elevated IOP, a better understanding of mechanisms that underlie IOP dysregulation in patients undergoing steroid treatments would be beneficial for prophylaxis and treatment. Our study supports the validity of mouse models to understand ocular mechanotransduction, brings insight into the molecular mechanism of corticosteroid-impaired pressure sensing and suggests a novel treatment that transcends current ciliary body/ciliary muscle-centered approaches. Specifically, ML-402 increased the permeability of TM monolayers (*20*), augmented conventional outflow (Fig. 2) and reversed DEX-induced IOP elevations, with a single eye drop sufficient to induce a 6-8 hour reduction to the normotensive baseline. Thus, progression of OHT in steroid glaucoma could reflect gradual dysregulation of the mechanosensory outflow setpoint that may be reversed through manipulation of mechanosensitive K^+^ efflux.

## Acknowledgements & Funding

We thank Dr. Daniel Minor (University of California of San Francisco) for the generous gifts of ML-402 (Lolicato et al., 2017).

The study was supported by the National Institutes of Health (T32EY024234 to CNR and DK, R01EY022076, R01EY031817, P30EY014800), Crandall Glaucoma Initiative, Stauss-Rankin Foundation, and an Unrestricted Grant from Research to Prevent Blindness to the Department of Ophthalmology at the University of Utah.

